# Cortical and spinal contributions to remote interlimb facilitation in humans

**DOI:** 10.64898/2026.05.02.722378

**Authors:** Atsushi Sasaki, Tatsuya Kato, Naotsugu Kaneko, Yohei Masugi, Matija Milosevic, Kimitaka Nakazawa

**Affiliations:** Graduate School of Arts and Sciences, The University of Tokyo, Tokyo, Japan; Sony Computer Science Laboratories, Inc., Tokyo, Japan; Japan Society for the Promotion of Science, Tokyo, Japan; School of Health Sciences, Tokyo International University, Saitama, Japan; The Miami Project to Cure Paralysis, University of Miami Miller School of Medicine, FL, USA; Department of Neurological Surgery, University of Miami Miller School of Medicine, FL, USA; Department of Biomedical Engineering, University of Miami, FL, USA

**Keywords:** remote effect, interlimb neural interaction, transcranial magnetic stimulation, intracortical inhibition, intracortical facilitation, F-wave

## Abstract

Voluntary contraction in one limb can facilitate motor output in a distant limb, a phenomenon commonly referred to as the remote effect. However, the neural mechanisms underlying this remote interlimb facilitation remain unclear. This study investigated cortical and spinal contributions to the remote effect in able-bodied participants. Transcranial magnetic stimulation (TMS) was applied over the hand area of the primary motor cortex using posterior–anterior (PA) and anterior–posterior (AP) current directions, which are sensitive to different cortical inputs. Cortical excitability was assessed using single- and paired-pulse paradigms to measure short-interval intracortical inhibition (SICI), short-interval intracortical facilitation (SICF), and short-latency afferent inhibition (SAI). Spinal motoneuron excitability was assessed from F-waves elicited by peripheral nerve stimulation. During voluntary lower-limb contractions, single-pulse TMS elicited larger motor evoked potentials in hand muscles across current directions, indicating a broad increase in net corticospinal output. However, only AP-sensitive paired-pulse measures showed reduced SICI and enhanced SICF during contraction, whereas PA-sensitive SICI and SICF were not significantly altered, suggesting that cortical modulation during the remote effect is expressed more clearly in AP-sensitive measures. SAI with PA stimulation was less consistently expressed during contraction, suggesting that afferent-related inhibitory modulation may also be influenced during the remote effect. In parallel, F-wave amplitude and persistence increased, consistent with enhanced spinal motoneuron excitability. Together, these results provide converging evidence that the remote effect in humans involves broad corticospinal and spinal facilitation, accompanied by current direction-dependent modulation of cortical excitability measures.

**KEY POINTS SUMMARY:**

- Voluntary contraction in one limb can facilitate motor output in a distant limb, but the mechanisms underlying this remote interlimb facilitation remain unclear.
- We tested whether remote lower-limb contraction modulates corticospinal output, intracortical excitability, and spinal motoneuron excitability in a resting hand muscle.
- Single-pulse transcranial magnetic stimulation showed that motor evoked potentials in the hand were facilitated during remote lower-limb contraction across multiple current directions, indicating a broad increase in net corticospinal output.
- Paired-pulse measures were modulated preferentially with anterior–posterior stimulation, with reduced short-interval intracortical inhibition and increased short-interval intracortical facilitation, suggesting current direction-dependent modulation of cortical excitability measures.
- F-wave amplitude and persistence were also enhanced during remote lower-limb contraction, indicating increased spinal motoneuron excitability. These findings provide converging evidence that the remote effect involves both cortical and spinal contributions.

## 1. INTRODUCTION

Voluntary movement of one limb can modulate motor output in a distant, non-engaged limb, a phenomenon commonly referred to as the “remote effect” (Tazoe and Komiyama, 2014). This form of remote interlimb facilitation provides a tractable model for investigating how neural activity in one limb influences corticospinal output to another. For instance, activating lower-limb muscles can facilitate corticospinal excitability in upper-limb muscles even when the target upper-limb muscle remains at rest, and vice versa (Tazoe and Komiyama, 2014). These findings suggest that upper- and lower-limb motor systems are functionally linked rather than entirely independent. The remote effect has been studied using transcranial magnetic stimulation (TMS), which measures corticospinal excitability via motor evoked potentials (MEPs) (Sasaki et al., 2021, 2020; Tazoe et al., 2007b). Although prior work shows that corticospinal output is modulated during remote limb activity, the circuit-level mechanisms underlying such modulation remain unclear. Understanding these mechanisms may help clarify a physiological substrate that contributes to broader forms of interlimb interaction during multi-limb motor behavior.

One approach to probing the mechanisms of remote interlimb facilitation is to use the sensitivity of TMS to the direction of induced current, which biases recruitment toward different inputs to corticospinal neurons in primary motor cortex (M1) (Day et al., 1989; Di Lazzaro et al., 2001). Posterior–anterior (PA) currents tend to evoke early indirect (I)-waves, which are highly synchronized with corticospinal output, whereas anterior–posterior (AP) currents preferentially recruit late I-waves, reflecting less synchronized and more polysynaptic cortical inputs (Aberra et al., 2020; Spampinato, 2020). Additionally, latero–medial (LM) currents are used to bias stimulation toward more direct activation of corticospinal axons, reducing dependence on intracortical processing (Werhahn et al., 1994). Thus, varying current direction can help determine whether remote interlimb facilitation preferentially affects specific cortical input streams and whether the observed changes are better explained by cortical versus downstream (subcortical/spinal) mechanisms.

Importantly, PA- and AP-sensitive responses appear to differ not only in the timing of I-wave recruitment but also in their sensitivity to task state and to inputs arising beyond M1, including premotor and cerebellar influences (Federico and Perez, 2017; Hamada et al., 2014; Volz et al., 2015). Because remote interlimb facilitation emerges during voluntary activation of a distant limb and is therefore likely to involve changes in distributed motor network state rather than only local corticospinal gain, AP-sensitive measures may be particularly susceptible to modulation. However, whether the remote effect preferentially modulates AP-sensitive cortical measures remains unknown.

Moreover, while single-pulse TMS studies have shown general facilitation of MEPs during remote limb activation, MEPs reflect the net summation of excitatory and inhibitory processes at cortical and spinal levels and cannot distinguish specific mechanisms. To address this, we designed a series of experiments to investigate how the remote effect modulates cortical and spinal excitability across complementary physiological measures. We combined single-pulse TMS across current directions with paired-pulse protocols focused on PA and AP orientations to assess short-interval intracortical inhibition (SICI), short-interval intracortical facilitation (SICF), and short-latency afferent inhibition (SAI)—well-established measures that probe intracortical inhibitory and excitatory mechanisms, linked primarily to GABA_A_-mediated inhibition (SICI), glutamatergic facilitation (SICF), and cholinergic mechanisms related to afferent inhibition (SAI), respectively (Lei and Perez, 2017; Ni et al., 2011; Rossini et al., 2015; Tokimura et al., 1996; Ziemann et al., 1998). To evaluate spinal contributions, we also assessed F-waves elicited by supramaximal peripheral nerve stimulation.

We hypothesized that remote lower-limb activity would modulate AP-sensitive measures more strongly than PA-sensitive measures, such that SICI would be reduced and SICF enhanced during remote contraction. This prediction was based on previous evidence that AP-sensitive responses are biased toward later I-wave-related inputs and can be modulated differently from PA-sensitive responses according to motor state and influences arising beyond M1 (Federico and Perez, 2017; Hamada et al., 2014; Volz et al., 2015). We also expected increased F-wave excitability, consistent with reports that remote voluntary contractions enhance spinal responsiveness (Kato et al., 2019; Masugi et al., 2019; Sasaki et al., 2020). This approach tests whether remote interlimb facilitation reflects a broad increase in corticospinal output, differential modulation of cortical excitability measures, and/or enhanced spinal motoneuron excitability. Clarifying these mechanisms may provide physiological insight into how activity in one limb influences motor output in another.

## 2. METHODS

### 2.1. Ethical approval

Written informed consent was obtained from all participants before participation, in accordance with the Declaration of Helsinki. All experimental procedures were approved by the Institutional Ethics Committee at the University of Tokyo (Approval No. 754).

### 2.2. Participants

None of the participants reported a history of neurological or musculoskeletal disorders. Exclusion criteria included any contraindication to TMS, including metal implants, cardiac pacemakers, a history of seizures or epilepsy, brain injury, neurosurgery, psychiatric illness, or the use of antidepressants or other neuromodulatory medications, in accordance with established safety guidelines (Rossi et al., 2021).

### 2.3. Electromyographic (EMG) activity

Surface EMG was recorded unilaterally from the right first dorsal interosseous (FDI) muscle of the dominant hand in all participants. Bipolar Ag/AgCl electrodes (Vitrode F-150S, Nihon Kohden, Tokyo, Japan) were placed over the muscle belly with an interelectrode distance of approximately 1 cm. A ground electrode was placed over the right olecranon. The skin was cleaned with alcohol to reduce skin impedance prior to electrode placement. Signals were band-pass filtered (5–1,000 Hz) and amplified (×1,000) using a multichannel amplifier (MEG-6108, Nihon Kohden, Tokyo, Japan). Data were digitized at a sampling frequency of 4,000 Hz through an analog-to-digital (A/D) converter (PowerLab/16SP, ADInstruments, Castle Hill, Australia) and stored on a computer for offline analysis.

### 2.4. Transcranial magnetic stimulation (TMS)

Experiments 1, 2, and 3 employed TMS over the left M1 using a monophasic magnetic stimulator (BiStim^2^, Magstim Co., Whitland, UK) with a figure-of-eight coil (loop diameter of 70 mm; Magstim Co., Whitland, UK). To bias recruitment toward different inputs to corticospinal neurons, the orientation of the induced current was varied across three configurations: (1) PA (coil handle at 45° to the midline), (2) AP (180° from PA), and (3) LM (handle perpendicular to the midline). These coil orientations are thought to preferentially recruit neural elements associated with early I-waves (PA), late I-waves (AP), and D-waves (LM), respectively (Di Lazzaro et al., 2012; Hamada et al., 2013; Sakai et al., 1997; Volz et al., 2015) (Figure 1A).

**Figure 1.**
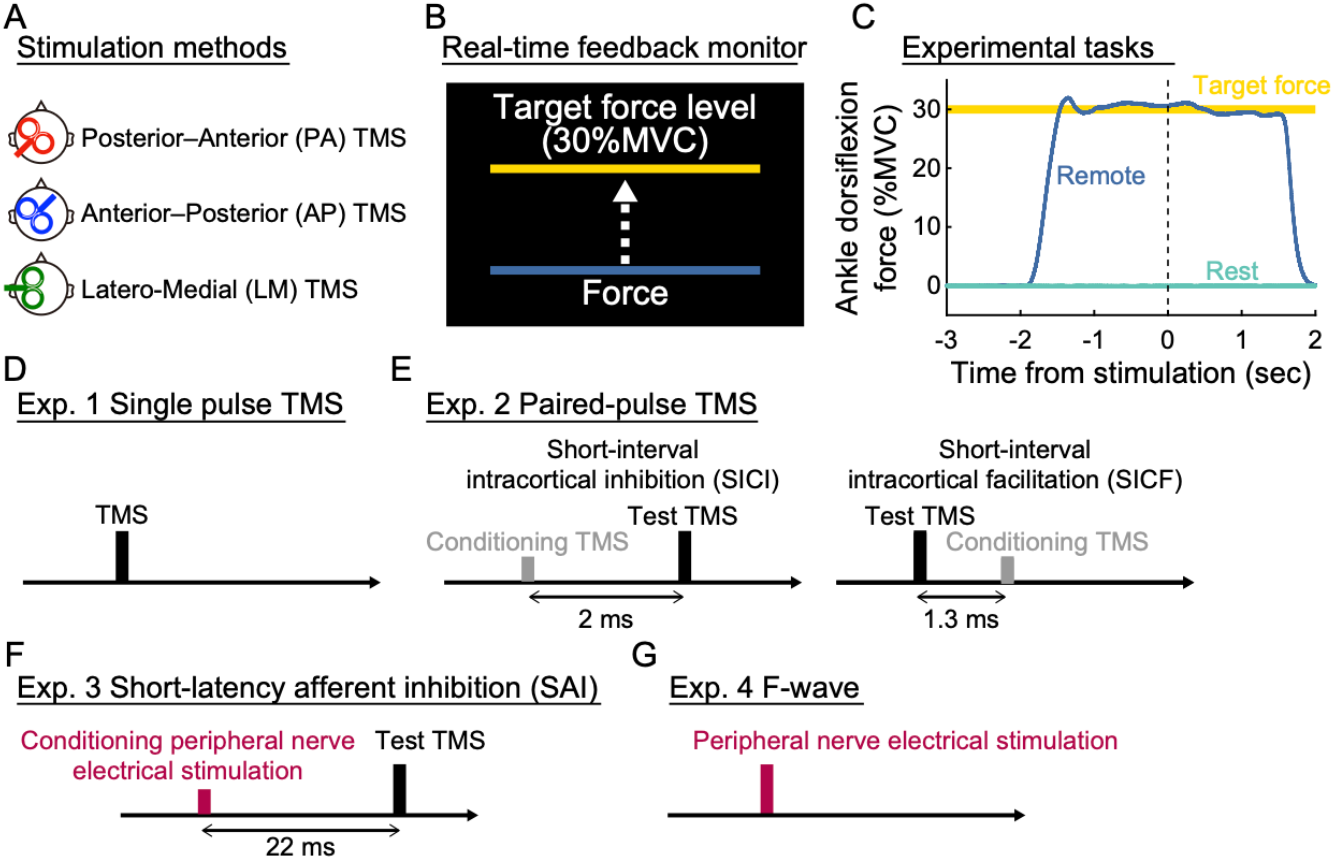
**(A)** Schematic of coil orientations for transcranial magnetic stimulation (TMS). Posterior– anterior (PA) and anterior–posterior (AP) currents are thought to bias recruitment toward different inputs to corticospinal neurons in primary motor cortex (M1), whereas latero–medial (LM) current is thought to favor more direct activation of corticospinal axons. **(B)** During the experimental task, participants received real-time visual feedback to maintain ankle dorsiflexion force at 30% of their maximum voluntary contraction (MVC). **(C)** Representative ankle dorsiflexion force traces during Rest and Remote conditions. In the Remote condition, participants maintained isometric dorsiflexion for approximately 4–5 seconds; in the Rest condition, stimulation was delivered without voluntary contraction. **(D)** Experiment 1 employed single-pulse TMS with PA, AP, or LM orientations to compare motor evoked responses elicited under different current directions. **(E)** Experiment 2 used paired-pulse TMS to assess short-interval intracortical inhibition (SICI; 2 ms interstimulus interval) and facilitation (SICF; 1.3 ms interstimulus interval). **(F)** In Experiment 3, short-latency afferent inhibition (SAI) was tested by applying peripheral electrical stimulation to the ulnar nerve 22 ms prior to a TMS pulse over M1. **(G)** In Experiment 4, spinal excitability was assessed by eliciting F-wave responses via supramaximal electrical stimulation of the ulnar nerve.

The optimal stimulation site (“hotspot”) for the FDI muscle was first identified using PA-oriented TMS. As previous studies suggest that coil orientation does not substantially alter hotspot location (Sakai et al., 1997), the same stimulation site was used for all orientations. Once the hotspot was identified, coil position and orientation were continuously monitored and maintained using a neuronavigation system (Brainsight, Rogue Research, Montreal, Canada). For the AP and LM conditions, the coil orientation was adjusted within the Brainsight system while maintaining the same cortical target.

All three orientations (PA, AP, and LM) were tested in Experiment 1. In Experiments 2 and 3, only PA and AP orientations were used (see Sections 2.5.2–2.5.4 for protocol details).

### 2.5. Experimental procedures

#### 2.5.1. General protocol

The study comprised four separate experiments. In each session, participants sat comfortably in an adjustable chair with their limbs supported to minimize movement. The right foot was securely positioned in a custom-made footplate with the ankle joint fixed at a neutral angle (90°) to ensure consistent biomechanical alignment across trials. After a brief warm-up and task familiarization, maximal voluntary contraction (MVC) of the right ankle dorsiflexors was assessed. Participants performed three maximal isometric dorsiflexion contractions, and force output was measured using a strain gauge sensor (LCB03K025L, A&D Company Limited, Japan) mounted to a fixed metal frame. The average of the three trials was used to determine each participant’s MVC.

During the experimental tasks, participants maintained a steady isometric dorsiflexion at 30% MVC, guided by real-time visual feedback displayed on a monitor (Figure 1B). TMS or peripheral nerve stimulation was delivered under two conditions: (1) at rest (Rest), and (2) during sustained isometric dorsiflexion at 30% MVC for approximately 4–5 seconds (Remote) (Figure 1C). In the Remote condition, stimulation was delivered after the verbal cue, once participants reached and maintained the target force level.

Throughout all sessions, participants were instructed to keep their hands and upper limbs fully relaxed. Surface EMG from the right FDI was continuously monitored to confirm the absence of overt involuntary activation. Offline EMG analysis further confirmed that background FDI activity did not differ significantly between conditions (see Section 3).

#### 2.5.2. Experiment 1: Single-pulse TMS

Fifteen healthy volunteers (5 females, 10 males; mean age ± SD: 28.1 ± 5.4 years) participated in Experiment 1. To investigate corticospinal responses elicited under different current directions during remote interlimb facilitation, MEPs were recorded from the right FDI muscle in response to single-pulse TMS delivered at three coil orientations: (1) PA, (2) AP, and (3) LM, under two task conditions: Rest and Remote (Figure 1D).

Stimulation intensity was individually adjusted to elicit MEPs of approximately 0.5 mV at rest for PA and AP orientations, and 1.0 mV for LM orientation, consistent with previous studies (Ibáñez et al., 2020; Ni et al., 2011; Sugawara et al., 2005). A higher intensity was used for LM to favor more direct activation of corticospinal axons, consistent with previous work (Werhahn et al., 1994). The mean stimulation intensities across participants were 58.9 ± 9.4% (PA), 74.3 ± 11.3% (AP), and 68.3 ± 7.9% (LM) of maximal stimulator output.

Each participant completed three coil orientation sessions (PA, AP, LM), with a minimum rest interval of 5 minutes between sessions. Within each session, Rest and Remote conditions were tested in a counterbalanced order, separated by at least 3 minutes. Each condition included 12 trials, with inter-trial intervals of approximately 10 seconds. The order of coil orientations and conditions was randomized across participants.

MEP onset latency was analyzed to verify that the three current directions produced the expected differences in response latency. The corresponding statistical results are reported in Section 3.1 and were consistent with previous work showing that PA, AP, and LM stimulation preferentially engage neural elements associated with early I-waves, late I-waves, and D-waves, respectively (Di Lazzaro et al., 2012; Hamada et al., 2013; Ni et al., 2011; Sakai et al., 1997; Volz et al., 2015).

#### 2.5.3. Experiment 2: Paired-pulse TMS

Twelve healthy adults (3 females, 9 males; mean age ± SD: 27.4 ± 3.1 years) participated in Experiment 2. To assess intracortical inhibition and facilitation during remote lower-limb activation, SICI and SICF were tested using established paired-pulse TMS paradigms (Kujirai et al., 1993; Long et al., 2017; Tokimura et al., 1996; Ziemann et al., 1998). Stimulation was delivered with the coil in PA and AP orientations during both Rest and Remote conditions (Figure 1E).

For SICI, the conditioning stimulus (CS) was set at 70–80% of the resting motor threshold (RMT) (PA: 71.7±3.7%; AP: 74.6± 4.7%) based on prior studies (Benavides et al., 2020; Chiou et al., 2013b; Long et al., 2017). For each current direction, stimulus intensities were defined relative to the RMT obtained with the same coil orientation. The test stimulus (TS) intensity was adjusted to evoke approximately 0.5 mV MEPs at rest (Chiou et al., 2013b). Because SICI magnitude depends on test MEP size (Long et al., 2017), the TS intensity in the Remote condition was adjusted so that test MEP amplitudes were matched to those in the Rest condition (i.e., approximately 0.5 mV) (Chiou et al., 2013b; Long et al., 2017). The interstimulus interval (ISI) was 2 ms (Long et al., 2017) (Figure 1E).

For SICF, the CS was set at 90–100% RMT (PA: 90.8±2.8%; AP: 90.9 ± 2.8%) (Cash et al., 2015; Long et al., 2017). The TS intensity was adjusted to elicit MEPs with an amplitude of approximately 0.5 mV at rest (Chiou et al., 2013b). Test MEP size during the Remote condition was adjusted to match MEP amplitudes produced at rest (i.e., approximately 0.5 mV) (Long et al., 2017). The CS was delivered 1.3 ms after the TS, at an ISI commonly used to assess SICF (Long et al., 2017) (Figure 1E).

Each participant completed two coil orientation sessions (PA and AP), separated by at least 5 minutes of rest. Within each session, 12 trials were recorded for each of the following: single-pulse TS alone (unconditioned MEP), SICI (CS + TS), and SICF (TS + CS), under both Rest and Remote conditions. Trials were separated by approximately 10 seconds, and each condition was separated by at least 3 minutes of rest. The order of coil orientations and stimulation paradigms was randomized across participants.

SICI and SICF were quantified as the ratio of conditioned to unconditioned MEP amplitude, expressed as a percentage: SICI and SICF (%) = (conditioned MEP / unconditioned test MEP) × 100 (Long et al., 2017).

#### 2.5.4. Experiment 3: TMS conditioned by peripheral electrical stimulation

Thirteen healthy adults (4 females, 9 males; mean age ± SD: 28.2 ± 5.8 years) participated in Experiment 3. To assess how PA- and AP-sensitive corticospinal responses are modulated by sensory afferent input, we employed a well-established SAI protocol, in which TMS over the left M1 was conditioned by peripheral electrical stimulation of the right ulnar nerve (Ni et al., 2011; Tokimura et al., 2000). Although both SAI and SICI are commonly interpreted as reflecting intracortical inhibitory processes, SAI engages cholinergic and GABA_A_ergic mechanisms involved in sensorimotor integration, distinct from the predominantly GABA_A_ergic inhibition underlying SICI (Rossini et al., 2015).

Peripheral conditioning stimuli were delivered to the right ulnar nerve at the wrist using a constant-current stimulator (DS7A; Digitimer Ltd., United Kingdom). Stimulation intensity was adjusted to elicit a visible thumb twitch (mean intensity: 10.7 ± 2.3 mA), corresponding to approximately three times the individual’s perception threshold (3.2 ± 0.7 mA), consistent with previous protocols (Ni et al., 2011).

As in Experiment 2, the TS was set to evoke approximately 0.5 mV MEPs at rest for each coil orientation (Cash et al., 2015). Given that Remote dorsiflexion increased MEP amplitudes in Experiment 1, TS intensities during the Remote condition were adjusted to match test MEP amplitudes obtained during Rest (i.e., approximately 0.5 mV) (Lei and Perez, 2017). The peripheral conditioning stimulus was applied 22 ms prior to the TMS pulse (Ni et al., 2011) (Figure 1F).

Each participant completed two coil orientation sessions (PA and AP), with at least 5 minutes of rest between sessions. Within each session, 12 trials were recorded for both unconditioned (TMS alone) and conditioned (peripheral stimulation + TMS) MEPs during both Rest and Remote conditions. Trials were separated by approximately 10 seconds, and at least 3 minutes of rest was provided between conditions. Session order was randomized across participants.

SAI was quantified as the percentage ratio of conditioned to unconditioned MEP amplitude: SAI (%) = (conditioned MEP / unconditioned test MEP) × 100 (Lei and Perez, 2017).

#### 2.5.5. Experiment 4: F-wave

Fourteen healthy adults (5 females, 9 males; mean age ± SD: 27.7 ± 5.6 years) participated in Experiment 4. To evaluate changes in spinal motoneuron excitability during the Remote condition, F-waves were elicited via supramaximal electrical stimulation of the right ulnar nerve at the wrist, following an established protocol (McNeil et al., 2013) (Figure 1G).

Stimulation was delivered using a constant-current stimulator (DS7A; Digitimer Ltd., UK) with a single monophasic square-wave pulse (200 μs duration). The cathode and anode electrodes were placed 3 cm apart, with the cathode positioned proximally. The stimulation intensity was gradually increased to identify the minimum current at which a maximal M-wave response (M_max_) was obtained, defined as the point beyond which further increases in current no longer increased M-wave amplitude. F-wave recordings were then acquired at 150% of M_max_ intensity, consistent with prior work (Long et al., 2017).

To isolate F-wave responses, a high-pass filter of 150 Hz was applied to ensure that the M-wave returned to baseline before F-wave onset (Tazoe and Perez, 2017).

Participants underwent both Rest and Remote conditions, with order randomized across individuals. In each condition, two blocks of 15 stimuli were delivered (30 stimuli total), with at least 3 min of rest between blocks and between conditions. F-wave amplitude and persistence served as outcome measures of spinal motoneuron excitability.

### 2.6. Data analysis

To quantify background EMG activity, the root mean square of the raw EMG signal was calculated within a 50-ms window immediately preceding each TMS or peripheral electrical stimulus using custom scripts written in MATLAB (R2024a, MathWorks, Natick, MA, USA). It is well established that MEPs elicited by TMS are facilitated by background muscle activation (Hallett, 2007). Therefore, any co-activation of the FDI during ankle dorsiflexion would confound interpretation of remote effect. To ensure that the FDI muscle remained inactive during the Remote condition, background EMG values were compared between the Rest and Remote conditions.

To quantify corticospinal modulation during the Remote condition, peak-to-peak MEP amplitudes from the FDI muscle were calculated for each trial across all experiments. In Experiment 1, MEP onset latency was analyzed to verify whether each TMS coil orientation produced response characteristics consistent with differential engagement of underlying neural pathways. Latency was determined by visual inspection with reference to a threshold corresponding to 2 standard deviations above the baseline mean computed over a 100-ms pre-stimulus period (Federico and Perez, 2017).

In Experiment 4, spinal excitability was assessed using F-wave parameters. F-wave amplitude was defined as the peak-to-peak value, and persistence was calculated as the percentage of trials (out of 30 stimuli per condition) in which F-wave peak-to-peak amplitude exceeded 50 μV (Rossini et al., 2015).

For all outcome measures (background EMG, MEP amplitude, MEP latency, F-wave amplitude, and persistence), values were averaged across trials within each condition and stimulation paradigm before statistical analysis.

### 2.7. Statistics

Data normality was assessed using the Shapiro–Wilk test. When the assumption of normality was violated, non-parametric alternatives were applied.

In Experiment 1, a one-way repeated-measures ANOVA was conducted to compare MEP onset latencies across the three coil orientations (PA, AP, and LM) during the Rest condition, to confirm that the three coil orientations produced different response latencies. When significant effects were observed, post hoc comparisons were conducted using paired t-tests with Bonferroni correction. Additionally, paired t-tests were used to compare MEP amplitude, onset latency, and background EMG between Rest and Remote conditions for each coil orientation.

In Experiments 2 and 3, SICI, SICF, and SAI were expressed as conditioned-to-unconditioned MEP ratios (%), where 100% represented no conditioning effect. For each coil orientation, Friedman tests were used to compare values across three levels: 100%, Rest, and Remote. When significant effects were detected, pairwise comparisons were performed using Wilcoxon signed-rank tests with Bonferroni correction. Background EMG activity was analyzed using the same approach.

In Experiment 4, Wilcoxon signed-rank tests were used to compare F-wave amplitude, F-wave persistence, and background EMG between Rest and Remote conditions.

All statistical analyses were performed using SPSS Statistics version 25 (IBM Corp., Armonk, NY, USA). Statistical significance was defined as p < 0.05. Effect sizes were reported using ηp^2^ for ANOVA, Kendall’s W for Friedman tests, Cohen’s d_z_ for paired t-tests, and r for Wilcoxon signed-rank tests.

## 3. RESULTS

### 3.1. Remote effect on corticospinal excitability (Experiment 1)

Figures 2A and 2B show representative average rectified MEP waveforms recorded during the Rest condition following TMS with PA, AP, and LM coil orientations in one participant. A one-way repeated-measures ANOVA revealed a significant effect of coil orientation on MEP onset latency (F(2,28) = 83.725, p < 0.001, ηp^2^ = 0.857). Post hoc pairwise comparisons indicated that onset latency was shortest with LM stimulation, longest with AP, and intermediate with PA, with all differences reaching statistical significance (Figure 2C; LM vs. PA: t(14) = 5.933, p < 0.001, d_z_ = 1.532; PA vs. AP: t(14) = 7.416, p < 0.001, d_z_ = 1.915; LM vs. AP: t(14) = 12.653, p < 0.001, d_z_ = 3.267). These results were consistent with differential corticospinal response characteristics across coil orientations.

**Figure 2.**
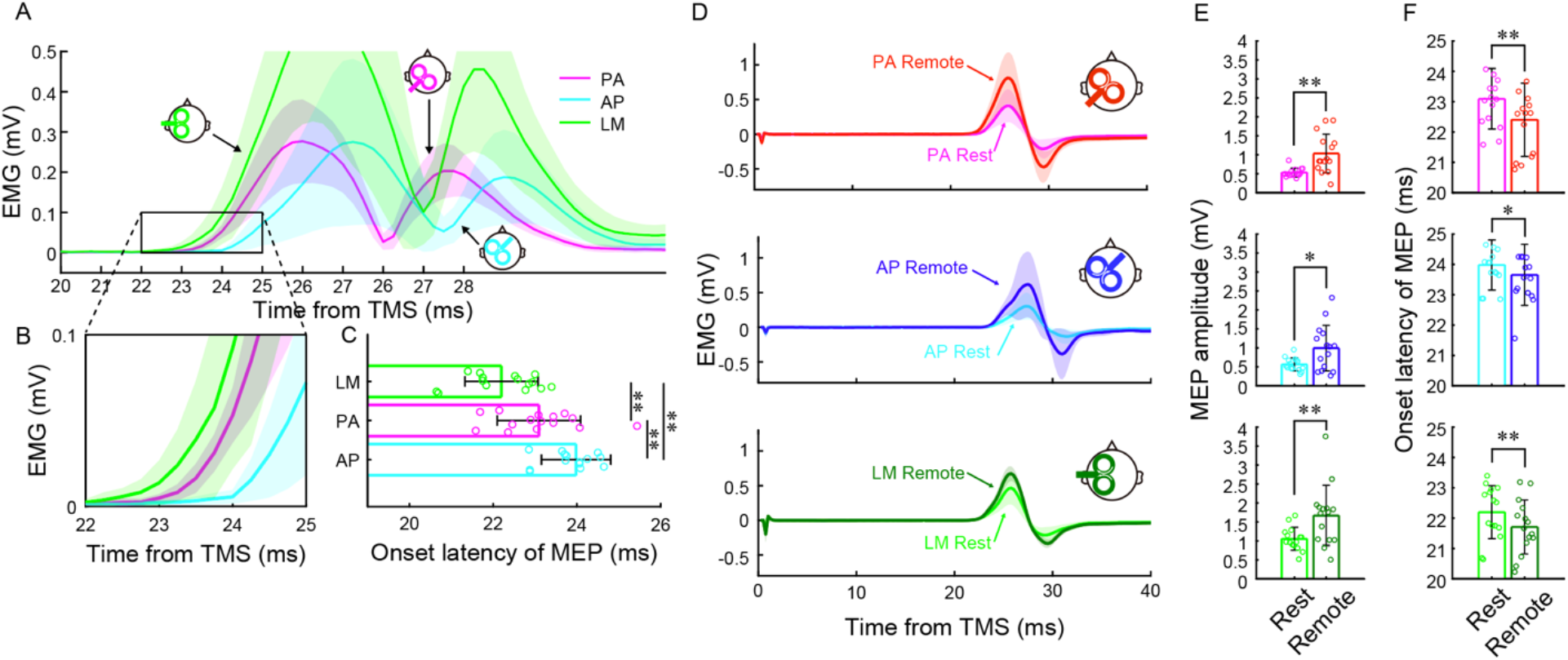
**(A)** Averaged rectified motor evoked potential (MEP) traces of the first dorsal interosseous (FDI) muscle following transcranial magnetic stimulation (TMS) with three coil orientations: posterior–anterior (PA), anterior–posterior (AP), and latero–medial (LM). Shaded areas indicate standard deviations. **(B)** Enlarged MEP traces illustrating differences in MEP onset latency across coil orientations. **(C)** Group data showing mean MEP onset latency during the Rest condition for PA, AP, and LM TMS. **(D)** Averaged MEP traces from a representative participant, showing Rest and Remote conditions for PA, AP, and LM TMS. **(E)** Group data showing mean MEP amplitudes during the Rest and Remote conditions for PA, AP, and LM TMS. **(F)** Group data showing mean MEP onset latency during the Rest and Remote conditions for PA, AP, and LM TMS. Error bars indicate standard deviations. *p < 0.05; **p < 0.01.

Figure 2D shows representative MEP traces from both Rest and Remote conditions for all three coil orientations. Paired t-tests revealed significantly larger MEP amplitudes in the Remote condition compared with Rest for PA (t(14) = -3.972, p = 0.001, d_z_ = -1.026), AP (t(14) = -2.779, p = 0.015, d_z_ = -0.718), and LM (t(14) = -3.182, p = 0.007, d_z_ = -0.822) stimulation (Figure 2E). Additionally, MEP onset latencies were significantly shorter in the Remote condition than in Rest for PA (t(14) = 6.047, p < 0.001, d_z_ = 1.561), AP (t(14) = 2.565, p = 0.022, d_z_ = 0.662), and LM (t(14) = 3.529, p = 0.003, d_z_ = 0.911) orientations (Figure 2F).

Importantly, background EMG activity in the FDI muscle did not significantly differ between Rest and Remote conditions for any coil orientation (PA: t(14) = -1.951, p = 0.071, d_z_ = -0.504; AP: t(14) = 0.007, p = 0.994, d_z_ = 0.002; LM: t(14) = -1.207, p = 0.247, d_z_ = -0.312), confirming that FDI muscle activation did not contribute to the observed MEP changes.

### 3.2. Remote effect on intracortical inhibition and facilitation (Experiment 2)

Figure 3A illustrates representative MEP waveforms from a single participant during the SICI protocol with PA stimulation. A Friedman test revealed a significant effect of condition on PA SICI ratio values [χ^2^(2) = 18.667, p < 0.001, W = 0.778] (Figure 3B). Post hoc Wilcoxon signed-rank tests showed that SICI ratio values were significantly lower than 100% (test MEP) in both the Rest (p = 0.007, r = 0.883) and Remote (p = 0.007, r = 0.883) conditions, with no significant difference between Rest and Remote (p = 1.000, r = 0.249).

**Figure 3.**
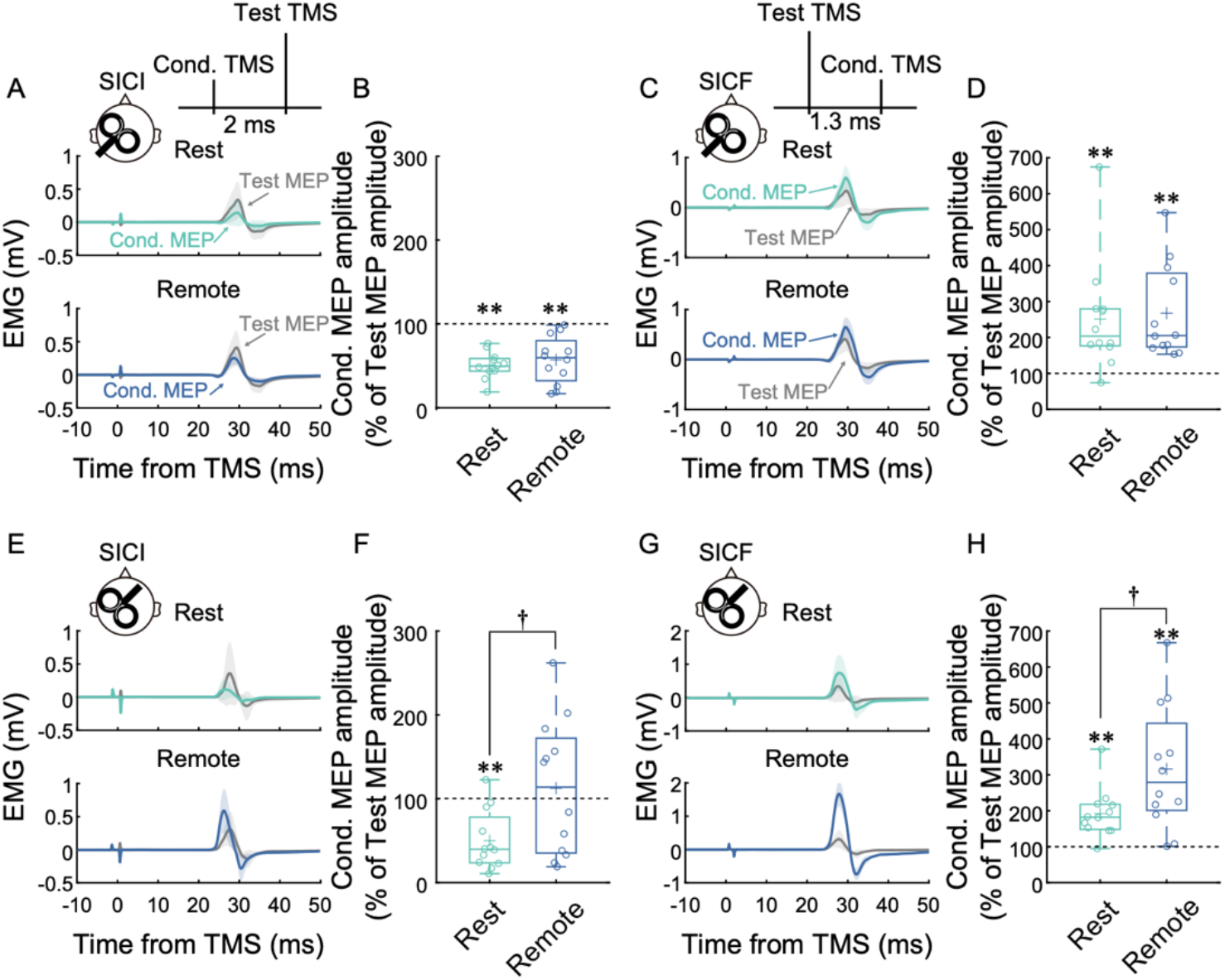
**(A)** Averaged motor evoked potential (MEP) traces of the first dorsal interosseous (FDI) muscle from a representative participant following transcranial magnetic stimulation (TMS) with posterior–anterior (PA) coil orientation during the Rest and Remote conditions in the short-interval intracortical inhibition (SICI) protocol. Shaded areas indicate standard deviations. **(B)** Boxplots showing group SICI ratio values (% of Test MEP amplitude) during the Rest and Remote conditions for PA stimulation. The dashed line indicates 100%, corresponding to no conditioning effect. **(C)** Averaged MEP traces of the FDI muscle from a representative participant following TMS with PA coil orientation during the Rest and Remote conditions in the short-interval intracortical facilitation (SICF) protocol. **(D)** Boxplots showing group SICF ratio values (% of Test MEP amplitude) during the Rest and Remote conditions for PA stimulation. **(E)** Averaged MEP traces of the FDI muscle from a representative participant following TMS with anterior–posterior (AP) coil orientation during the Rest and Remote conditions in the SICI protocol. **(F)** Boxplots showing group SICI ratio values (% of Test MEP amplitude) during the Rest and Remote conditions for AP stimulation. **(G)** Averaged MEP traces of the FDI muscle from a representative participant following TMS with AP coil orientation during the Rest and Remote conditions in the SICF protocol. **(H)** Boxplots showing group SICF ratio values (% of Test MEP amplitude) during the Rest and Remote conditions for AP stimulation. Boxes represent the 25th and 75th percentiles, and whiskers indicate the minimum and maximum values. Asterisks indicate significant differences from the no-effect value of 100% (Test MEP). Daggers indicate significant differences between the Rest and Remote conditions. **p < 0.01; ^†^p < 0.05.

Figure 3C shows representative MEP waveforms for SICF with PA stimulation. A Friedman test indicated a significant effect of condition on PA SICF ratio values [χ^2^(2) = 15.167, p < 0.001, W = 0.632] (Figure 3D). Post hoc analyses revealed that SICF ratio values were significantly greater than 100% in both the Rest (p = 0.009, r = 0.608) and Remote (p = 0.007, r = 0.624) conditions, with no significant difference between Rest and Remote (p = 1.000, r = 0.128).

There were no significant differences in background EMG activity across conditions for the PA SICI [χ^2^(3) = 1.000, p = 0.801, W = 0.028] and SICF [χ^2^(3) = 3.700, p = 0.296, W = 0.103] protocols.

Figure 3E illustrates representative MEP traces obtained using AP stimulation in the SICI protocol. A Friedman test revealed a significant effect of condition on AP SICI ratio values [χ^2^(2) = 8.167, p = 0.017, W = 0.340] (Figure 3F). Post hoc analysis showed that SICI ratio values were significantly lower than 100% (test MEP) in the Rest condition (p = 0.014, r = 0.815), but not in the Remote condition (p = 1.000, r = 0.136). Moreover, SICI ratio values were significantly greater in the Remote than in the Rest condition (p = 0.029, r = 0.747).

Figure 3G shows representative MEP traces for AP stimulation during the SICF protocol. A Friedman test revealed a significant effect of condition on AP SICF ratio values [χ^2^(2) = 18.500, p < 0.001, W = 0.771] (Figure 3H). Post hoc tests demonstrated that SICF ratio values were significantly greater than 100% in both the Rest (p = 0.009, r = 0.861) and Remote (p = 0.007, r = 0.883) conditions. SICF ratio values were also significantly greater in the Remote condition than in the Rest condition (p = 0.029, r = 0.747).

No significant differences were observed in background EMG activity across conditions in the AP SICI protocol [χ^2^(3) = 3.900, p = 0.272, W = 0.108]. Although the Friedman test indicated a condition effect on background EMG in the AP SICF protocol [χ^2^(3) = 8.100, p = 0.044, W = 0.225], post hoc comparisons did not reveal any significant differences (all p > 0.100).

### 3.3. Remote effect on afferent inhibition (Experiment 3)

Figure 4A shows representative MEP traces recorded during the Rest and Remote conditions in the SAI protocol using PA coil orientation. A Friedman test revealed a significant effect of condition on PA SAI ratio values [χ^2^(2) = 15.846, p < 0.001, W = 0.609] (Figure 4B). Post hoc Wilcoxon signed-rank tests showed that SAI ratio values were significantly lower than 100% (test MEP) in the Rest condition (p = 0.004, r = 0.882), but not in the Remote condition (p = 0.057, r = 0.649). There was no significant difference in SAI ratio values between Rest and Remote conditions (p = 0.588, r = 0.359). Background EMG activity did not differ significantly across conditions [χ^2^(3) = 6.323, p = 0.097, W = 0.162].

**Figure 4.**
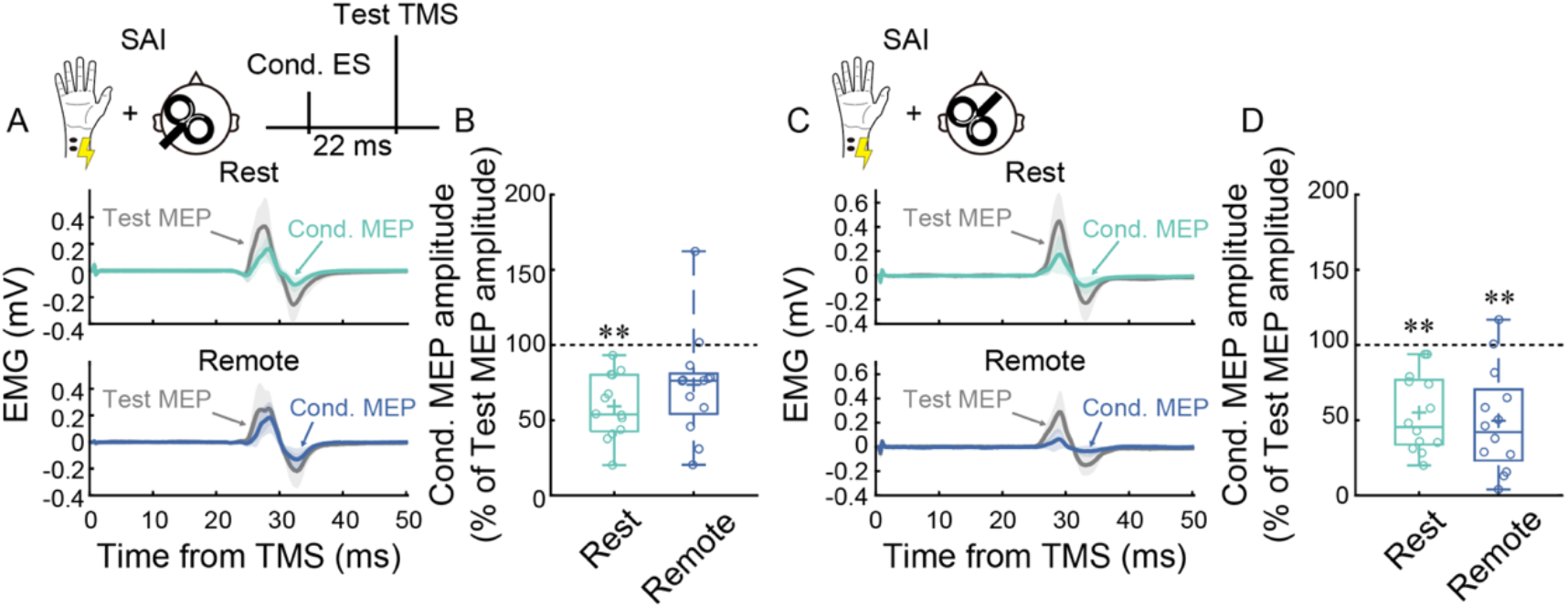
**(A)** Averaged motor evoked potential (MEP) traces of the first dorsal interosseous (FDI) muscle from a representative participant following transcranial magnetic stimulation (TMS) with posterior–anterior (PA) coil orientation during the Rest and Remote conditions in the short-latency afferent inhibition (SAI) protocol. Shaded areas indicate standard deviations. **(B)** Boxplots showing group PA SAI ratio values (% of Test MEP amplitude) during the Rest and Remote conditions. **(C)** Averaged MEP traces of the FDI muscle from a representative participant following TMS with anterior–posterior (AP) coil orientation during the Rest and Remote conditions in the SAI protocol. **(D)** Boxplots showing group AP SAI ratio values (% of Test MEP amplitude) during the Rest and Remote conditions. The dashed line indicates 100%, corresponding to no conditioning effect. Boxes represent the 25th and 75th percentiles and whiskers indicate the minimum and maximum values. Asterisks indicate significant differences from the no-effect value of 100%. **p < 0.01.

Figure 4C shows MEP traces recorded under the same protocol using AP coil orientation. A Friedman test revealed a significant effect of condition on AP SAI ratio values [χ^2^(2) = 14.000, p < 0.001, W = 0.538] (Figure 4D). Post hoc analysis showed that SAI ratio values were significantly lower than 100% in both the Rest (p = 0.004, r = 0.882) and Remote (p = 0.009, r = 0.824) conditions. However, no significant difference was found between Rest and Remote conditions (p = 1.000, r = 0.145). Background EMG activity did not differ significantly across conditions [χ^2^(3) = 4.569, p = 0.206, W = 0.117].

### 3.4. Remote effect on spinal excitability (Experiment 4)

Figure 5A illustrates representative M- and F-wave traces evoked by supramaximal ulnar nerve stimulation during the Rest and Remote conditions. F-wave amplitude and persistence were significantly increased during the Remote condition compared with Rest (amplitude: p < 0.001, r = 0.881; persistence: p = 0.001, r = 0.865) (Figure 5B). Background EMG activity in the FDI did not differ significantly between conditions (p = 0.177, r = 0.361), confirming that the observed changes in spinal excitability were not attributable to involuntary hand muscle activation.

**Figure 5.**
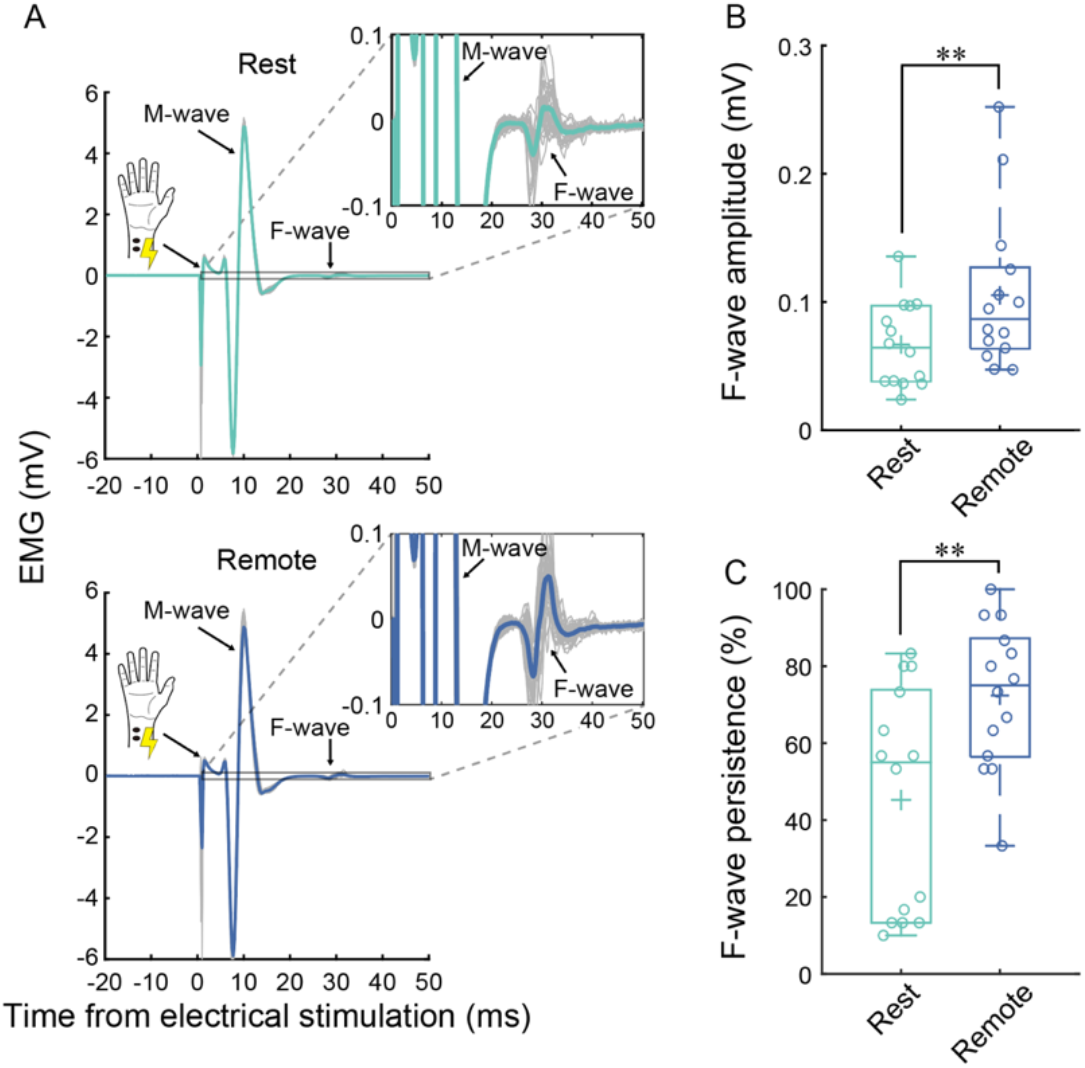
**(A)** Averaged M- and F-wave responses recorded from the first dorsal interosseous (FDI) muscle in a representative participant following ulnar nerve stimulation during the Rest and Remote conditions. Thick-colored lines indicate average waveforms, whereas thin gray lines indicate individual trials. **(B)** Boxplots showing group data for F-wave amplitude (mV) during the Rest and Remote conditions. **(C)** Boxplots showing group data for F-wave persistence (%) during the Rest and Remote conditions. Boxes represent the 25th and 75th percentiles and whiskers indicate the minimum and maximum values. Asterisks indicate significant differences between Rest and Remote conditions. ** p < 0.01.

## 4. DISCUSSION

This study investigated how voluntary lower-limb contraction modulates corticospinal and spinal excitability in a remote upper-limb muscle. Using a remote effect paradigm, we combined single-pulse TMS delivered with different current directions, paired-pulse TMS measures of intracortical excitability, and F-wave recordings to assess converging cortical and spinal contributions. Across current directions, remote contraction increased MEP amplitude and shortened MEP onset latency (Experiment 1), indicating a robust increase in net corticospinal output. Notably, intracortical physiology was modulated in a current direction-dependent manner: AP-sensitive measures showed reduced SICI and enhanced SICF during remote contraction (Experiment 2), whereas PA-sensitive SICI and SICF did not show comparable changes. In addition, SAI was less consistently expressed with PA stimulation during remote contraction (Experiment 3), and F-wave amplitude and persistence were increased (Experiment 4), supporting a downstream contribution at the level of spinal motoneurons.

These findings suggest that the remote effect involves both cortical and spinal contributions, with the cortical component expressed in a current direction-dependent manner. In particular, the paired-pulse findings indicate that remote contraction does not modulate intracortical excitability uniformly, but instead alters the inhibitory-facilitatory balance more clearly in AP-sensitive measures. This pattern is broadly consistent with our initial hypothesis that AP-sensitive measures associated with late I-waves would be preferentially modulated during remote limb contraction. The differential modulation across PA- and AP-sensitive measures suggests that the interlimb remote effect is accompanied by non-uniform modulation of intracortical processes.

### 4.1. Cortical and spinal contributions to the remote effect

TMS current direction is commonly used to bias recruitment toward different inputs to corticospinal neurons. PA currents tend to evoke early I-waves, whereas AP currents preferentially recruit late I-waves, which are thought to reflect less synchronized and more polysynaptic cortical inputs (Di Lazzaro and Ziemann, 2013). LM orientation is commonly used to bias stimulation toward more direct activation of corticospinal axons, thereby reducing dependence on intracortical processing (Werhahn et al., 1994). Within this framework, Experiment 1 demonstrated that remote lower-limb contraction facilitated MEPs and shortened MEP onset latency across PA, AP, and LM orientations. This suggests that the remote effect enhances net corticospinal output via mechanisms that are not limited to a single intracortical input stream.

The facilitation of LM-evoked MEPs is consistent with increased excitability downstream of intracortical processing, potentially at subcortical and/or spinal levels. This interpretation is further supported by Experiment 4, where F-wave amplitude and persistence, indices of spinal motoneuron excitability, were significantly enhanced during the Remote condition. These findings align with prior reports showing enhanced spinal excitability during lower-limb activation (Kato et al., 2019; Masugi et al., 2019; Sasaki et al., 2020) and collectively indicate that spinal circuitry is an integral component of remote effect.

At the same time, the paired-pulse findings indicate that the cortical contribution is not expressed uniformly across current directions. Experiment 2 showed that remote contraction reduced SICI and enhanced SICF with AP stimulation, whereas comparable changes were not observed with PA stimulation. Because SICI and SICF are commonly interpreted as reflecting inhibitory and facilitatory intracortical processes, respectively, this dissociation suggests that the remote effect is accompanied by non-uniform modulation of intracortical excitability across current directions. These measures likely involve GABA_A_-mediated inhibition and glutamatergic excitation, respectively (Di Lazzaro et al., 2007; Rossini et al., 2015; Ziemann et al., 2015, 1996), implicating these neurotransmitter systems in interlimb remote facilitation. In contrast, SICI and SICF elicited by PA stimulation were unchanged despite the facilitation of PA single-pulse MEPs during the Remote condition. This dissociation suggests that PA-sensitive corticospinal facilitation during the remote effect may reflect different mechanisms not well captured by these paired-pulse measures alone, including changes in other cortical processes and/or increased gain at subcortical or spinal stages.

Experiment 3 further suggests that afferent-related modulation may contribute differentially across current directions. With PA stimulation, SAI was robust at rest but appeared less consistently expressed during remote contraction. Given that SAI depends on cholinergic mechanisms implicated in sensorimotor integration (Di Lazzaro et al., 2005, 2000), these findings, combined with prior reports of reduced cortical silent periods during remote effect paradigms (Tazoe et al., 2007b, 2007a), point to a potential role for non-GABAergic inhibitory systems in PA-sensitive remote facilitation. Together, these findings suggest that PA-driven remote facilitation may rely more on downstream gain changes or afferent-driven modulation than on the inhibitory–facilitatory balance captured by SICI and SICF. This interpretation should remain cautious, however, as the direct Rest–Remote contrast did not reach significance and SAI likely reflects a composite of multiple mechanisms.

The current findings also refine prior work by Chiou et al. (2013a) who reported reduced PA-sensitive SICI during contralateral lower-limb contractions. In contrast, we did not observe significant PA SICI modulation during the ipsilateral remote contraction used here. This discrepancy may reflect differences in the pathways supporting ipsilateral versus contralateral remote interlimb contraction. For example, contralateral remote effects could involve transcallosal contributions, whereas ipsilateral effects likely engage intrahemispheric sensorimotor networks. Future work directly comparing these conditions could further elucidate the underlying circuitry.

### 4.2. Why were AP-sensitive measures modulated more clearly during the remote effect?

A key question raised by our findings is why AP-sensitive measures showed clearer modulation than PA-sensitive measures during the remote effect. One plausible explanation is that AP-sensitive responses are biased toward later I-wave-related inputs and are more strongly influenced by task state and by inputs arising beyond M1, including premotor and cerebellar influences (Federico and Perez, 2017; Hamada et al., 2014; Volz et al., 2015). In this context, the present findings suggest that AP-sensitive paired-pulse measures may be particularly informative for detecting the cortical modulation associated with remote contraction, which may involve a change in overall motor network state rather than only local corticospinal gain. This interpretation is consistent with previous work showing that AP-sensitive responses are particularly sensitive to changes in motor state, including movement preparation (Hannah et al., 2018; Ibáñez et al., 2020).

A further consideration is that directional TMS does not provide a one-to-one mapping onto single inhibitory or facilitatory pathways. Previous work has shown that AP- and PA-sensitive measures can reflect partially distinct but overlapping physiological processes, and that the circuits recruited by AP stimulation are influenced not only by current direction but also by pulse parameters such as pulse duration (Fong et al., 2021; Hannah and Rothwell, 2017). Because pulse duration was not manipulated in the present study, the AP-sensitive effects observed here should not be interpreted as evidence for selective modulation of a single intracortical inhibitory or facilitatory circuit. Instead, they are more appropriately interpreted as current direction-dependent differences in the modulation captured by the physiological measures used to probe cortical excitability during the remote effect.

Broader network influences, including cerebellar contributions, should also be considered. Prior studies have shown that cerebellar-M1 interactions differ across PA- and AP-sensitive responses and can vary with behavioral state and learning (Hamada et al., 2014; Spampinato et al., 2020). This raises the possibility that the present AP-sensitive effects reflect not only local cortical changes within M1 but also broader network influences engaged during remote contraction. Because the present study did not assess cerebellar-M1 connectivity directly or manipulate cerebellar excitability, the contribution of specific non-primary motor areas cannot be determined. However, the present findings suggest that the enhancement of corticospinal output during the remote effect may reflect influences from a broader motor network rather than changes confined to M1 alone.

### 4.3. Limitations

Several limitations should be acknowledged. First, we examined a single interlimb configuration, in which voluntary ankle dorsiflexion modulated corticospinal output to an intrinsic hand muscle (FDI). Remote effects are context dependent, and both the magnitude and direction of modulation can vary with the muscles engaged and the behavioral demands of the task (Sasaki et al., 2021). Accordingly, it remains uncertain whether the current direction-dependent pattern observed here, clear modulation of AP-sensitive intracortical measures alongside more global facilitation of corticospinal and spinal output, generalizes to other muscle pairings (e.g., upper-to-lower limb interactions), more proximal effectors, or different contraction modes (e.g., dynamic or bilateral tasks).

Second, cortical and spinal measures were obtained in separate experiments rather than within the same participants. Thus, although the present findings provide converging evidence that both cortical and spinal mechanisms contribute to the remote effect, they do not establish how these processes covary within individuals. In addition, although different TMS current directions are useful for biasing recruitment toward different corticospinal inputs, they do not isolate completely distinct neural pathways. Accordingly, the present findings should be interpreted as evidence for current direction-dependent modulation of cortical excitability measures, rather than as proof of selective recruitment of a single discrete intracortical circuit.

Nevertheless, the present results provide a controlled physiological characterization of the remote effect using complementary cortical and spinal measures. Combining multiple TMS measures (SICI, SICF, SAI) with a spinal index (F-waves) allowed us to characterize the remote effect at several levels of the motor system, while limiting variability introduced by heterogeneous task demands. Future studies should apply the same framework across multiple muscle pairings, bidirectional interlimb configurations, and different contraction intensities, and should also test whether the present physiological effects relate to behavior involving multiple effectors.

### 4.4. Conclusions

In summary, our results show that the remote effect is associated with increased corticospinal output and enhanced spinal motoneuron excitability during voluntary lower-limb contraction. Although single-pulse MEP facilitation was observed across all tested current directions, paired-pulse measures revealed clearer modulation in AP-sensitive measures than in PA-sensitive measures, indicating that the cortical component of the remote effect is expressed in a current direction-dependent manner. Importantly, the AP-sensitive effects observed here should not be interpreted as evidence for a mechanism confined to M1 alone. Rather, they raise the possibility that activity in a broader motor network, including regions outside primary motor cortex, contributes to the modulation of corticospinal output during the remote effect. In parallel, increased F-wave amplitude and persistence support a downstream contribution at the level of the spinal motoneuron pool. Together, these findings suggest that the remote effect reflects converging cortical and spinal contributions.

## ADDITIONAL INFORMATION

### Data availability statement

The data that support the findings of this study are available on request from the corresponding author.

### Competing interest

No conflicts of interest, financial or otherwise, are declared by the authors.

### Author contributions

A.S., T.K., N.K., and K.N. conceived and designed the research. A.S., T.K., and N.K. performed the experiments. A.S. analyzed the data. All authors interpreted the results of the experiments. A.S. drafted the manuscript. A.S., T.K., N.K., Y.M., M.M., and K.N. critically revised the manuscript for important intellectual content. All authors have read and approved the final version of this manuscript and agree to be accountable for all aspects of the work in ensuring that questions related to the accuracy or integrity of any part of the work are appropriately investigated and resolved. All persons designated as authors qualify for authorship, and all those who qualify for authorship are listed.

### Funding

This project was supported by a Grant-in-Aid for Early-Career Scientists (KAKENHI) from the Japan Society for the Promotion of Science (JSPS) to A.S. (23K16640).

## Notes

### Competing Interest Statement

The authors have declared no competing interest.

## REFERENCES

Aberra, A.S., Wang, B., Grill, W.M., Peterchev, A.V., 2020. Simulation of transcranial magnetic stimulation in head model with morphologically-realistic cortical neurons. Brain Stimulation 13, 175–189. 10.1016/j.brs.2019.10.002

Benavides, F.D., Jo, H.J., Lundell, H., Edgerton, V.R., Gerasimenko, Y., Perez, M.A., 2020. Cortical and Subcortical Effects of Transcutaneous Spinal Cord Stimulation in Humans with Tetraplegia. J. Neurosci. 40, 2633–2643. 10.1523/JNEUROSCI.2374-19.2020

Cash, R.F.H., Isayama, R., Gunraj, C.A., Ni, Z., Chen, R., 2015. The influence of sensory afferent input on local motor cortical excitatory circuitry in humans: Effects of sensory input on SICF. J Physiol 593, 1667–1684. 10.1113/jphysiol.2014.286245

Chiou, S.-Y., Wang, R.-Y., Liao, K.-K., Wu, Y.-T., Lu, C.-F., Yang, Y.-R., 2013a. Co-activation of primary motor cortex ipsilateral to muscles contracting in a unilateral motor task. Clinical Neurophysiology 124, 1353–1363. 10.1016/j.clinph.2013.02.001

Chiou, S.-Y., Wang, R.-Y., Liao, K.-K., Yang, Y.-R., 2013b. Homologous Muscle Contraction during Unilateral Movement Does Not Show a Dominant Effect on Leg Representation of the Ipsilateral Primary Motor Cortex. PLoS ONE 8, e72231. 10.1371/journal.pone.0072231

Day, B.L., Dressler, D., Maertens de Noordhout, A., Marsden, C.D., Nakashima, K., Rothwell, J.C., Thompson, P.D., 1989. Electric and magnetic stimulation of human motor cortex: surface EMG and single motor unit responses. The Journal of Physiology 412, 449–473. 10.1113/jphysiol.1989.sp017626

Di Lazzaro, V., Oliviero, A., Profice, P., Pennisi, M.A., Di Giovanni, S., Zito, G., Tonali, P., Rothwell, J.C., 2000. Muscarinic receptor blockade has differential effects on the excitability of intracortical circuits in the human motor cortex. Experimental Brain Research 135, 455–461. 10.1007/s002210000543

Di Lazzaro, V., Oliviero, A., Saturno, E., Pilato, F., Insola, A., Mazzone, P., Profice, P., Tonali, P., Rothwell, J., 2001. The effect on corticospinal volleys of reversing the direction of current induced in the motor cortex by transcranial magnetic stimulation. Experimental Brain Research 138, 268–273. 10.1007/s002210100722

Di Lazzaro, V., Pilato, F., Dileone, M., Profice, P., Ranieri, F., Ricci, V., Bria, P., Tonali, P.A., Ziemann, U., 2007. Segregating two inhibitory circuits in human motor cortex at the level of GABAA receptor subtypes: A TMS study. Clinical Neurophysiology 118, 2207–2214. 10.1016/j.clinph.2007.07.005

Di Lazzaro, V., Pilato, F., Dileone, M., Tonali, P.A., Ziemann, U., 2005. Dissociated effects of diazepam and lorazepam on short-latency afferent inhibition: Dissociated effects of benzodiazepines on SAI. The Journal of Physiology 569, 315–323. 10.1113/jphysiol.2005.092155

Di Lazzaro, V., Profice, P., Ranieri, F., Capone, F., Dileone, M., Oliviero, A., Pilato, F., 2012. I-wave origin and modulation. Brain Stimulation 5, 512–525. 10.1016/j.brs.2011.07.008

Di Lazzaro, V., Ziemann, U., 2013. The contribution of transcranial magnetic stimulation in the functional evaluation of microcircuits in human motor cortex. Front. Neural Circuits 7. 10.3389/fncir.2013.00018

Federico, P., Perez, M.A., 2017. Distinct Corticocortical Contributions to Human Precision and Power Grip. Cerebral Cortex 27, 5070–5082. 10.1093/cercor/bhw291

Fong, P.-Y., Spampinato, D., Rocchi, L., Hannah, R., Teng, Y., Di Santo, A., Shoura, M., Bhatia, K., Rothwell, J.C., 2021. Two forms of short-interval intracortical inhibition in human motor cortex. Brain Stimulation 14, 1340–1352. 10.1016/j.brs.2021.08.022

Hallett, M., 2007. Transcranial Magnetic Stimulation: A Primer. Neuron 55, 187–199. 10.1016/j.neuron.2007.06.026

Hamada, M., Galea, J.M., Di Lazzaro, V., Mazzone, P., Ziemann, U., Rothwell, J.C., 2014. Two Distinct Interneuron Circuits in Human Motor Cortex Are Linked to Different Subsets of Physiological and Behavioral Plasticity. Journal of Neuroscience 34, 12837–12849. 10.1523/JNEUROSCI.1960-14.2014

Hamada, M., Murase, N., Hasan, A., Balaratnam, M., Rothwell, J.C., 2013. The Role of Interneuron Networks in Driving Human Motor Cortical Plasticity. Cerebral Cortex 23, 1593–1605. 10.1093/cercor/bhs147

Hannah, R., Cavanagh, S.E., Tremblay, S., Simeoni, S., Rothwell, J.C., 2018. Selective Suppression of Local Interneuron Circuits in Human Motor Cortex Contributes to Movement Preparation. J. Neurosci. 38, 1264–1276. 10.1523/JNEUROSCI.2869-17.2017

Hannah, R., Rothwell, J.C., 2017. Pulse Duration as Well as Current Direction Determines the Specificity of Transcranial Magnetic Stimulation of Motor Cortex during Contraction. Brain Stimulation 10, 106–115. 10.1016/j.brs.2016.09.008

Ibáñez, J., Fu, L., Rocchi, L., Spanoudakis, M., Spampinato, D., Farina, D., Rothwell, J.C., 2020. Plasticity induced by pairing brain stimulation with motor-related states only targets a subset of cortical neurones. Brain Stimulation 13, 464–466. 10.1016/j.brs.2019.12.014

Kato, T., Sasaki, A., Yokoyama, H., Milosevic, M., Nakazawa, K., 2019. Effects of neuromuscular electrical stimulation and voluntary commands on the spinal reflex excitability of remote limb muscles. Exp Brain Res 237, 3195–3205. 10.1007/s00221-019-05660-6

Kujirai, T., Caramia, M.D., Rothwell, J.C., Day, B.L., Thompson, P.D., Ferbert, A., Wroe, S., Asselman, P., Marsden, C.D., 1993. Corticocortical inhibition in human motor cortex. The Journal of Physiology 471, 501–519. 10.1113/jphysiol.1993.sp019912

Lei, Y., Perez, M.A., 2017. Cortical contributions to sensory gating in the ipsilateral somatosensory cortex during voluntary activity: Sensory gating in the iS1. J Physiol 595, 6203–6217. 10.1113/JP274504

Long, J., Federico, P., Perez, M.A., 2017. A novel cortical target to enhance hand motor output in humans with spinal cord injury. Brain 140, 1619–1632. 10.1093/brain/awx102

Masugi, Y., Sasaki, A., Kaneko, N., Nakazawa, K., 2019. Remote muscle contraction enhances spinal reflexes in multiple lower-limb muscles elicited by transcutaneous spinal cord stimulation. Exp Brain Res 237, 1793–1803. 10.1007/s00221-019-05536-9

McNeil, C.J., Butler, J.E., Taylor, J.L., Gandevia, S.C., 2013. Testing the excitability of human motoneurons. Front. Hum. Neurosci. 7. 10.3389/fnhum.2013.00152

Ni, Z., Charab, S., Gunraj, C., Nelson, A.J., Udupa, K., Yeh, I.-J., Chen, R., 2011. Transcranial Magnetic Stimulation in Different Current Directions Activates Separate Cortical Circuits. Journal of Neurophysiology 105, 749–756. 10.1152/jn.00640.2010

Rossi, S., Antal, A., Bestmann, S., Bikson, M., Brewer, C., Brockmöller, J., Carpenter, L.L., Cincotta, M., Chen, R., Daskalakis, J.D., Di Lazzaro, V., Fox, M.D., George, M.S., Gilbert, D., Kimiskidis, V.K., Koch, G., Ilmoniemi, R.J., Lefaucheur, J.P., Leocani, L., Lisanby, S.H., Miniussi, C., Padberg, F., Pascual-Leone, A., Paulus, W., Peterchev, A.V., Quartarone, A., Rotenberg, A., Rothwell, J., Rossini, P.M., Santarnecchi, E., Shafi, M.M., Siebner, H.R., Ugawa, Y., Wassermann, E.M., Zangen, A., Ziemann, U., Hallett, M., 2021. Safety and recommendations for TMS use in healthy subjects and patient populations, with updates on training, ethical and regulatory issues: Expert Guidelines. Clinical Neurophysiology 132, 269–306. 10.1016/j.clinph.2020.10.003

Rossini, P.M., Burke, D., Chen, R., Cohen, L.G., Daskalakis, Z., Di Iorio, R., Di Lazzaro, V., Ferreri, F., Fitzgerald, P.B., George, M.S., Hallett, M., Lefaucheur, J.P., Langguth, B., Matsumoto, H., Miniussi, C., Nitsche, M.A., Pascual-Leone, A., Paulus, W., Rossi, S., Rothwell, J.C., Siebner, H.R., Ugawa, Y., Walsh, V., Ziemann, U., 2015. Non-invasive electrical and magnetic stimulation of the brain, spinal cord, roots and peripheral nerves: Basic principles and procedures for routine clinical and research application. An updated report from an I.F.C.N. Committee. Clinical Neurophysiology 126, 1071–1107. 10.1016/j.clinph.2015.02.001

Sakai, K., Ugawa, Y., Terao, Y., Hanajima, R., Furubayashi, T., Kanazawa, I., 1997. Preferential activation of different I waves by transcranial magnetic stimulation with a figure-of-eight-shaped coil. Exp Brain Res 113, 24–32. 10.1007/BF02454139

Sasaki, A., Kaneko, N., Masugi, Y., Kato, T., Milosevic, M., Nakazawa, K., 2021. Task- and Intensity-Dependent Modulation of Arm-Trunk Neural Interactions in the Corticospinal Pathway in Humans. eNeuro 8, ENEURO.0111-21.2021. 10.1523/ENEURO.0111-21.2021

Sasaki, A., Kaneko, N., Masugi, Y., Milosevic, M., Nakazawa, K., 2020. Interlimb neural interactions in corticospinal and spinal reflex circuits during preparation and execution of isometric elbow flexion. Journal of Neurophysiology 124, 652–667. 10.1152/jn.00705.2019

Spampinato, D., 2020. Dissecting two distinct interneuronal networks in M1 with transcranial magnetic stimulation. Exp Brain Res 238, 1693–1700. 10.1007/s00221-020-05875-y

Spampinato, D.A., Celnik, P.A., Rothwell, J.C., 2020. Cerebellar–Motor Cortex Connectivity: One or Two Different Networks? J. Neurosci. 40, 4230–4239. 10.1523/JNEUROSCI.2397-19.2020

Sugawara, K., Furubayashi, T., Takahashi, M., Ni, Z., Ugawa, Y., Kasai, T., 2005. Remote effects of voluntary teeth clenching on excitability changes of the human hand motor area. Neuroscience Letters 377, 25–30. 10.1016/j.neulet.2004.11.059

Tazoe, T., Endoh, T., Nakajima, T., Sakamoto, M., Komiyama, T., 2007a. Disinhibition of upper limb motor area by voluntary contraction of the lower limb muscle. Exp Brain Res 177, 419–430. 10.1007/s00221-006-0686-1

Tazoe, T., Komiyama, T., 2014. Interlimb neural interactions in the corticospinal pathways. JPFSM 3, 181–190. 10.7600/jpfsm.3.181

Tazoe, T., Perez, M.A., 2017. Cortical and reticular contributions to human precision and power grip: Control of precision and power grip. J Physiol 595, 2715–2730. 10.1113/JP273679

Tazoe, T., Sakamoto, M., Nakajima, T., Endoh, T., Komiyama, T., 2007b. Effects of remote muscle contraction on transcranial magnetic stimulation-induced motor evoked potentials and silent periods in humans. Clinical Neurophysiology 118, 1204–1212. 10.1016/j.clinph.2007.03.005

Tokimura, H., Lazzaro, V., Tokimura, Y., Oliviero, A., Profice, P., Insola, A., Mazzone, P., Tonali, P., Rothwell, J.C., 2000. Short latency inhibition of human hand motor cortex by somatosensory input from the hand. The Journal of Physiology 523, 503–513. 10.1111/j.1469-7793.2000.t01-1-00503.x

Tokimura, H., Ridding, M.C., Tokimura, Y., Amassian, V.E., Rothwell, J.C., 1996. Short latency facilitation between pairs of threshold magnetic stimuli applied to human motor cortex. Electroencephalography and Clinical Neurophysiology/Electromyography and Motor Control 101, 263–272. 10.1016/0924-980X(96)95664-7

Volz, L.J., Hamada, M., Rothwell, J.C., Grefkes, C., 2015. What Makes the Muscle Twitch: Motor System Connectivity and TMS-Induced Activity. Cereb. Cortex 25, 2346–2353. 10.1093/cercor/bhu032

Werhahn, K.J., Fong, J.K.Y., Meyer, B.U., Priori, A., Rothwell, J.C., Day, B.L., Thompson, P.D., 1994. The effect of magnetic coil orientation on the latency of surface EMG and single motor unit responses in the first dorsal interosseous muscle. Electroencephalography and Clinical Neurophysiology/ Evoked Potentials 93, 138–146. 10.1016/0168-5597(94)90077-9

Ziemann, U., Lönnecker, S., Steinhoff, B.J., Paulus, W., 1996. Effects of antiepileptic drugs on motor cortex excitability in humans: A transcranial magnetic stimulation study: Antiepileptic Drugs and Excitability of Human Cortex. Ann Neurol. 40, 367–378. 10.1002/ana.410400306

Ziemann, U., Reis, J., Schwenkreis, P., Rosanova, M., Strafella, A., Badawy, R., Müller-Dahlhaus, F., 2015. TMS and drugs revisited 2014. Clinical Neurophysiology 126, 1847–1868. 10.1016/j.clinph.2014.08.028

Ziemann, U., Tergau, F., Wassermann, E.M., Wischer, S., Hildebrandt, J., Paulus, W., 1998. Demonstration of facilitatory I wave interaction in the human motor cortex by paired transcranial magnetic stimulation. The Journal of Physiology 511, 181–190. 10.1111/j.1469-7793.1998.181bi.x

